# An investigation towards understanding the role of CDH1 gene in different types of Breast Cancers

**DOI:** 10.1101/2021.01.11.426236

**Authors:** Antara Sengupta, Raja Banerjee

## Abstract

At recent age breast cancer attracts the attention of both the medical and the scientific community for its deadly occurrence throughout the globe as it is considered to be happened due to genetic aberration. Out of several genes expressed, it is found that cadherin 1, type 1 (CDH1) is responsible in several ways to control the metabolic order in human. Hence we focus on CDH1 gene whether any deviation in it especially alteration/modification in its sequence is responsible for the occurrence of this deadly disease. Towards this end study of the available genomic sequences of CDH1 gene of several patients, suffering from various types of breast cancer (obtained from the Sanger Database), are studied. The results emphasizes that alternation/modification in the sequence of the CDH1 gene affect its regular function which may have a potential role in damaging the different types of breast tissues, causing malfunction and leading to breast cancers in patients.

## 1. Introduction

Cancer, the deadly disease, is caused by uncontrolled division of abnormal cells in a part of the body. As the cells undergo rapid proliferation, the disease spreads so rapidly and can become invasive. Moreover, cancers in different parts of the body affect the patients differently and also seek treatments in different ways. Researchers from almost all the fields accept the challenge of this almost incurable disease to overcome and prevent its worldwide fatal effect. In woman, breast cancer is the second leading cause of cancer related death that undergoes physical and mental trauma (http://ghr.nlm.nih.gov/gene/CDH1). It is increasing particularly in the developing countries where the majority of cases are diagnosed in late stages and hence, seeks intensive attention. Breast cancer can take place in different areas of the breast — the ducts, the lobules, or in some cases, the tissue in between. Till date from the histopathological point of view breast cancers are broadly categories as five following types: 1) Ductal Carcinoma (DC), 2) Lobular Carcinoma (LC), 3) Lobular Carcinoma *In Situ* (LCIS), 4) Ductal Carcinoma *In Situ* (DCIS), and 5) Ducto Lobular Carcinoma *In Situ* (DLCIS).

At recent age breast cancer draws the attention of the medical as well as the scientific community for its deadly occurrence and considered to be happened due to genetic aberration [1]. Literature survey shows [2] that different genes are responsible for maintaining different crucial biological functions and aberration of those are causing different types of breast cancers due to non-proper functioning. Out of several genes expressed, it is found that “cadherin 1, type 1” (CDH1) is responsible in several ways to control the metabolic order in human. Hence we focus on CDH1 gene whether any deviation in it especially alteration/modification in its sequence is responsible for the occurrence of this deadly disease. Hence, a detailed study of CDH1 gene from patients suffering from various types of breast cancer would be worthy as that can help to get consensus how deviation or alteration/modification in its sequence is responsible for the occurrence of this deadly disease. CDH1 belongs to the gene families CD (Cluster of Differentiation) and CDH (cadherins) and it’s chromosomal location is 16q22.1 [3]. The gene contains information for making a protein called ‘epithelial cadherin or E-cadherin’, which helps in cell adhesion, transmitting chemical signals within the cells, controlling cell maturation and movement, along with regulation of the activities of certain genes. It also acts as a tumor suppressor protein [4]. Mutations in CDH1 genes can be responsible for lack of the protein E-cadherin, which leads cells to grow too rapidly and divide in an uncontrolled way. It is seen that CDH1 is one of the most differentially expressed genes between ductal and lobular carcinomas at the mRNA level [4, 5]. According to a group of researchers, the loss of E-cadherin function, and the dysfunctional cell adhesion and regulation of acting cytoskeleton in Invasive lobular carcinoma (ILC) have been suggested to be the factor responsible for its distinct morphology, pattern of invasion, and metastatic behavior [2, 6, 7]. Whereas, some others have found that Loss of heterozygosity for chromosomes 16q (LOH at 16q) is the second most frequent somatic genetic event in breast cancer [1]. E-cadherin can be used as diagnostic marker for breast cancer [6]. Multiple studies of gene expression profiling have advanced the understanding of the molecular diagnosis of breast cancer, providing the background for oncologists to use the triple-negative phenotype to describe the basal-like molecular sub-type [8–13]. Numerous studies have been taken place to investigate and understand the characteristics of mutated CDH1 in different types of breast cancers [6–10]. It is worth to state that although reported work have tried to unfold facts and findings of the gene associated to particular type of breast cancer, till date no report has been found to focus on the quantitative analysis and comparison of the role of CDH1, playing crucial role for almost all the types of breast cancers. Here in this manuscript towards identifying the role of CDH1 regarding its driving force for the occurrence of breast cancer, we have investigated and analyzed the existing data set of breast cancer from the Sanger database, to get a patient wise comprehensive idea on the alteration/modification in their respective genomic sequence. This would provide an idea how the alteration in the genomic sequence plays crucial role to damage the normal function of the cells and causing breast cancer. Further, from the respective sequence analysis, a consensus can be drawn for causing the respective type of breast cancer as the synthesis of defective protein, due to alteration of primary sequence, would participate in malfunctioning of the system through changes in their physico-chemical behavior. The investigation would help to understand the perspective of the formation of several breast cancers.

## 2. Methods and Materials

### 2.1 Data Set Specification

To carry out the investigation related to the mutations of CDH1 gene in different patients, the data related to mutations have been taken from the database www.cancer.sanger.ac.uk. The total data set taken, are shown in the Table S1. The summary of the dataset specification is given in Table 1.

**Table 1.**
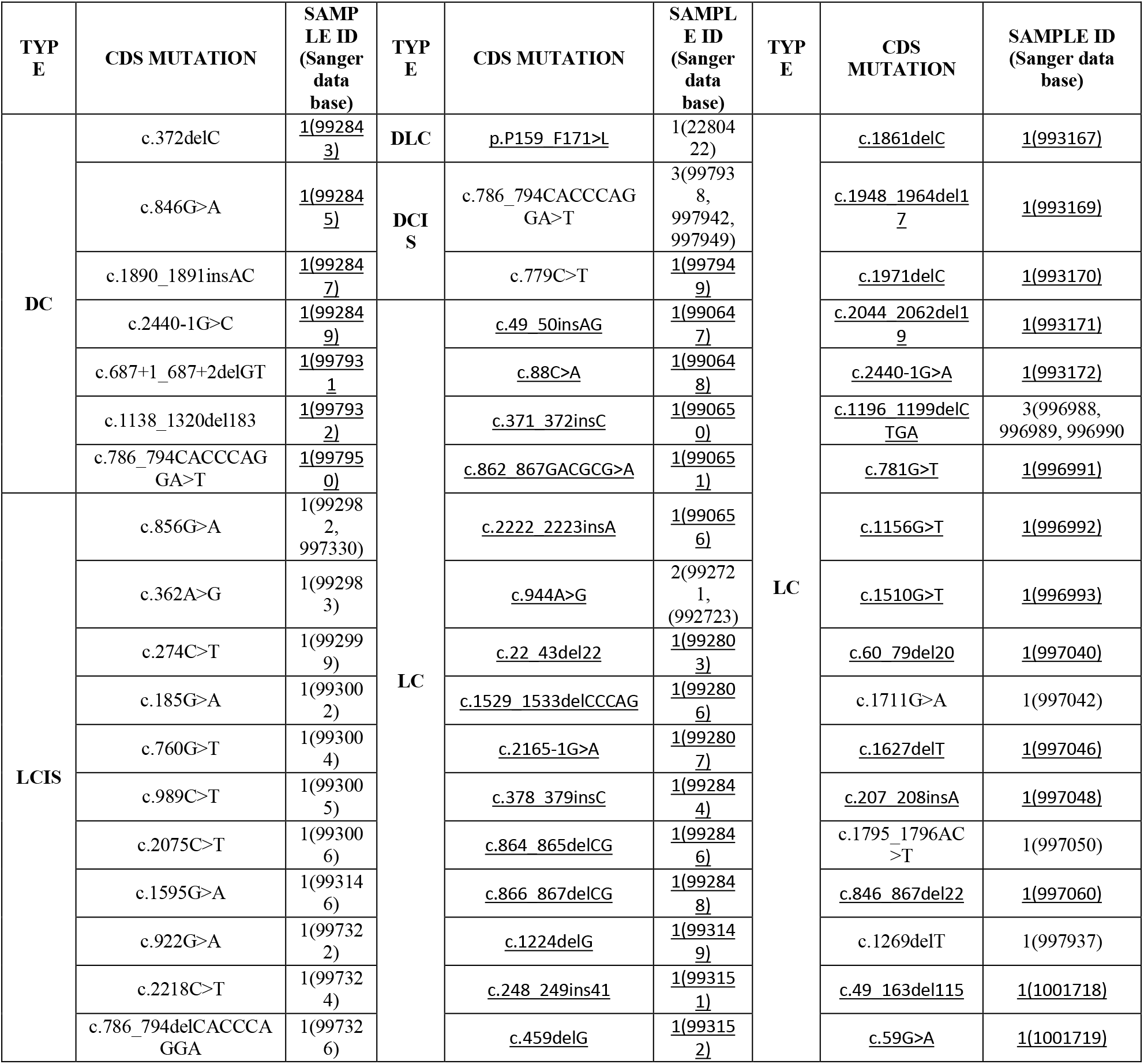
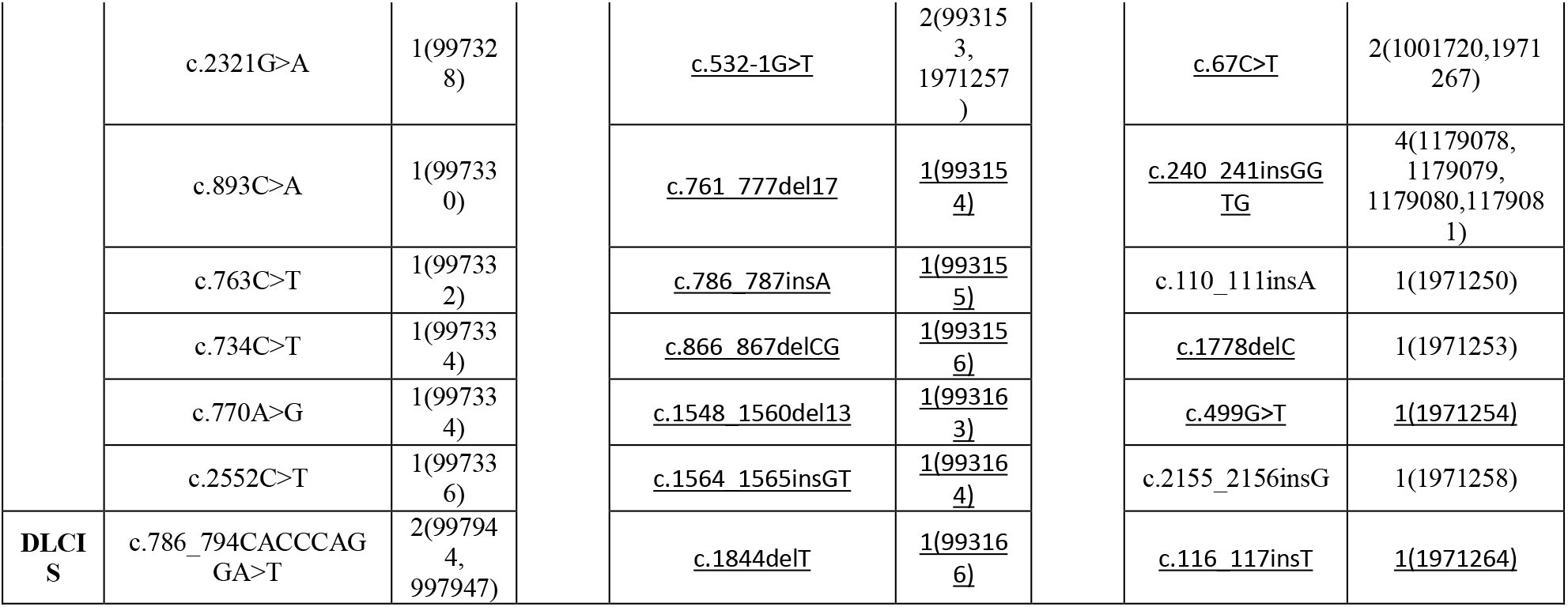

### 2.2 Methods

In this present work it is aimed to make quantitative understanding of the trend of mutations occurring in CDH1 gene which leads to occur breast cancer and to investigate the role of CDH1 in different types of breast cancers. Information related to mutations reported in table 1 is used to prepare mutant DNA sequences. The investigations have been taken place in three levels, i.e. DNA sequential level, primary protein sequential level and in the physico-chemical characteristics of mutated proteins.

#### 2.2.1 Preparing mutant data as reported in database

Normal, disease free (wild type) nucleotide sequence of CDH1 gene has been taken from NCBI database as the control. To prepare the mutant DNA sequence, mutation location has been identified (using inputs from Table S1) and the corresponding change(s) has been made accordingly as it is reported in Sanger Database. Thus total 86 mutated DNA sequences are constructed using customized program in Matlab as per patient data specification in Sanger database, which are shown in Table S1. Further, the mutated data set are translated into their corresponding amino acid sequences using online software https://www.expasy.org/ to have some biophysical characterization.

## 3. Result and Discussion

### 3.1 Investigating Patient dataset at DNA level

Mutations may be silent or missense. Substitution of a single nucleotide can make huge biological changes and be responsible for a disease. In this part of the manuscript it is tried to investigate and understand the trend of mutations occurred in DNA sequential level and finally become responsible for breast cancer at particular location.

#### 3.1.1 Investigating trends of mutations at DNA level

A keen analysis of the sequences obtained from the dataset for the patients as per information shown in Table S1, it is found that different type of mutations e.g. single point mutation, frame shift and other different kinds of changes have been acquired in the specific positions of the wild type CDH1 DNA sequence corresponding to patients. In Table 2 using pair-wise alignment of the wild type CDH1 DNA sequence and that obtained from the patient database as mentioned in Table S1, a short glimpse of the changes are highlighted which shade light on how the deviation/ mutation of the wild type nucleotide sequence would act as the cause of different kind of breast cancer. A detailed analysis have been done in subsequent parts to get a consensus if a type of breast cancer (e.g. DC, LC, DCIS etc) can be correlated with the type of mutation. A keen observation of Table 2, identifies even similar type of mutation (e.g. single point mutation / frame shift / insertion etc) at different positions in the wild type would cause different types of breast cancer. As an example it can be noted that substitution of similar G > A at position 846 causes DC while at position 856 causes LCIS.

**Table 2:**
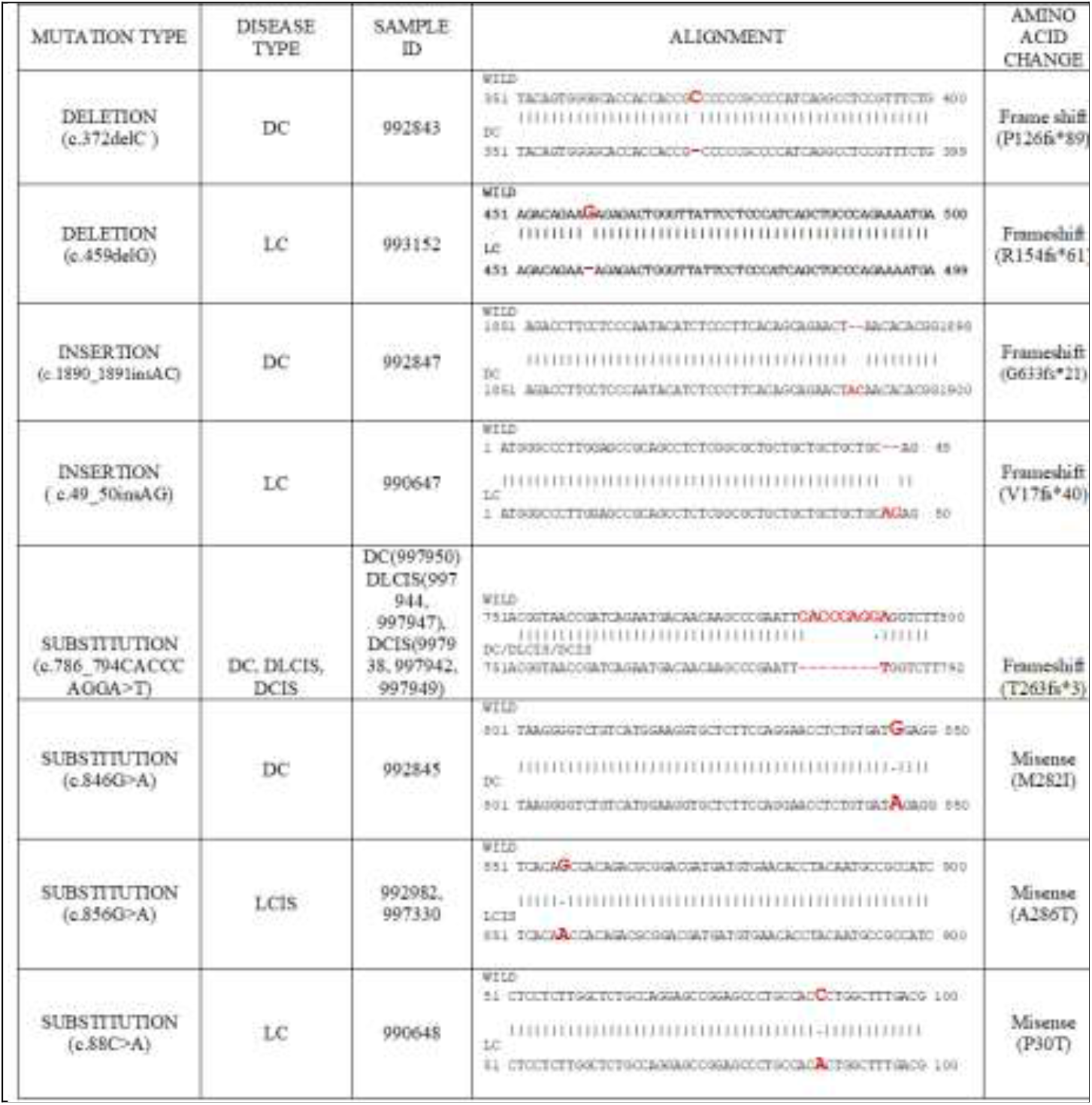
Pair-wise Alignment to identify the mutation location

Further, considering different types of mutations as observed, a percentage of each types taken place in 86 samples are calculated. It is identified that the mutations are majorly taken place due to different kinds of deletion frame shifts(28%), insertion frame shifts(18%) and complex frame shifts(9%) throughout all six types of breast cancer sequences. It can be easily observed from the Figure 1 that maximum numbers of frame shifts are caused due to deletion type of mutations. Further, different types of substitution mutation like missense (26%), intronic (23%) and nonsense (13%) also play crucial role. Finer observations towards patients’ data reveal more minute facts. Although deletion frame shift has the leading role in case of Lobular Carcinoma in Situ (LCIS) type of breast cancers, the mutations majorly have been occurred in the lobules or milk glands in the breast due to missense mutations and the change of a single base pair caused the substitution of a different amino acid in the resulting protein, whereas, in case of DC, intronic mutations have significant role.

**Fig 1:**
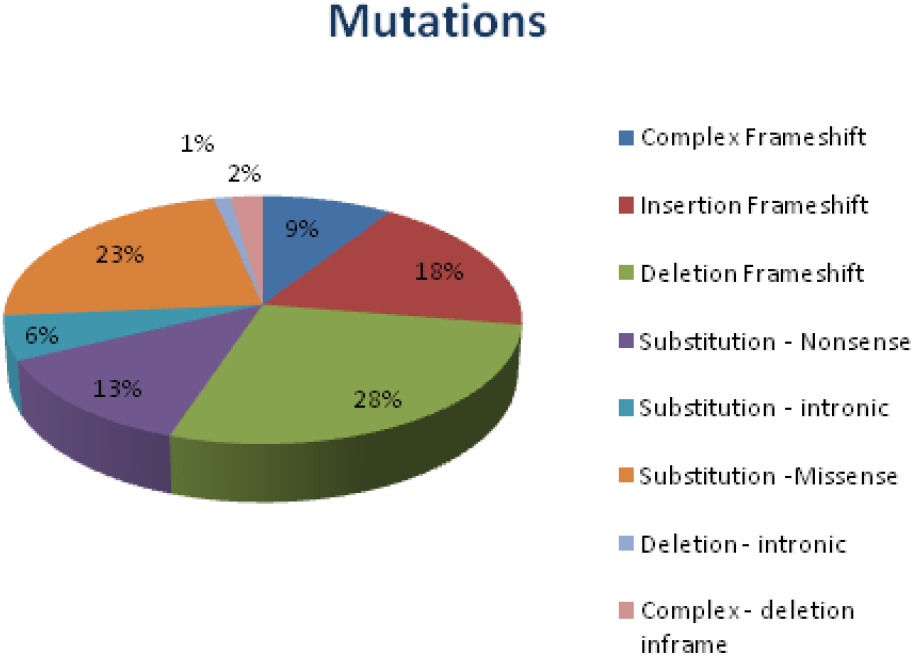
Different types of mutations with respect to total data set.

Although previously we have observed that similar type of mutation (e.g. substitution etc) located at different position of the wild type sequence causing different types of breast cancer, it is rather important to note that deletion frame shift occurred at 786^th^ position of CDS area (*786_794CACCCAGGA*) are found in four (DC, DCIS, DLCIS and LCIS) out of six types of breast cancers (shown in Table 3). However, point to be noted that out of this four, three are occurring in the duct region of the breast. Numerous biological reasons may there be responsible behind the fact and one of the vital reasons might be variations of Protein - Protein Interactions (PPI) of the mature proteins. However, this strongly emphasizes that the sequence *786_794CACCCAGGA* must have a definite importance in maintaining regular function of the system, otherwise cause fatal disease like breast cancer.

**Table 3:** Same mutations at different physical location of breast may cause different kinds of breast cancer

| TYPE | SAMPLE_ID | MUTATION |
| --- | --- | --- |
| DC | 997950 | c.786_794CACCCAGGA>T |
| DCIS | 997938 | c.786_794CACCCAGGA>T |
| DLCIS | 997944 | c.786_794CACCCAGGA>T |
| LCIS | 997326 | c.786_794delCACCCAGGA |

Several comparative analysis of mutations reveal that mutation rate is not same throughout a DNA sequence. Some coding locations or areas of the DNA sequence are more mutation prone. In the present work we have identified that LC type of cancer has significance with respect to locational importance (Table 4). The CDS areas c.532_1, c.67, c.786_787, c.1196_1199, c.944, c.866_867, c.240_241 are mutation prone and responsible for LC to occur.

**Table 4.** Mutation prone coding locations.

| TYPE | SAMPLE_ID | MUTATION | TYPE OF MUTATION |
| --- | --- | --- | --- |
| LC | <u>993153</u> | <u>c.532-1G>T</u> | Single point mutation |
| LC | <u>1971257</u> |  |  |
| LC | <u>1001720</u> | <u>c.67C>T</u> |  |
| LC | <u>1971267</u> |  |  |
| LC | <u>992721</u> | <u>c.944A>G</u> |  |
| LC | <u>992723</u> |  |  |
| LC | <u>1179078</u> | <u>c.240_241insXGGTG</u> | Insertion |
| LC | <u>1179079</u> |  |  |
| LC | <u>1179080</u> |  |  |
| LC | <u>1179081</u> |  |  |
| LC | <u>992848</u> | <u>c.866_867delCG</u> | Deletion |
| LC | <u>993156</u> |  |  |
| LC | <u>996988</u> | <u>c.1196_1199delCTGA</u> |  |
| LC | <u>996989</u> |  |  |
| LC | <u>996990</u> |  |  |
| LC | <u>993155</u> | <u>c.786_787insAInsertion</u> |  |

#### 3.1.2 Quantification of Bio-physical Properties of Mutated Nucleotides

Analysis of the patient wise data obtained from the Sanger Database as shown in previous subsection clearly reveals that different types of mutations at the nucleotides remains responsible for the different type of breast cancers. This ignites the thrust to get an insight whether the changes took place in nucleotide sequences can make any substantial changes in their different parameters of biophysical properties (e.g molecular weight, percentage of GC content, thermodynamic relationship between entropy (ΔS), enthalpy (ΔH), free energy (ΔG) etc) as the information stored in the nucleotide sequences translated to the expressed proteins and the modification of behavior of the single molecule can even influence the biological function of the organism. OligoCalc [14] web interface which can calculate the biophysical properties of oligonucelotides, is used to get quantitative understanding of the biophysical properties of CDH1(s) obtained due to different mutations and how does they vary in different types of breast cancers. According to the result derived it is observed that although the GC content remains unaltered, all the other properties have been changed due to mutation. The range of values for each parameter has also been differed from types of breast cancer. The changes of biophysical properties for each variant are shown in supplementary Table S3. A summary of the same shown in table 5 shed light on the minimum and maximum range of changes in comparison to wild type for each type of breast cancers.

**Table 5:** Quantification of Bio-physical Properties

| Type | MOL WT | GC COUNT | SALT ADJUSTED | BASIC | DELTA H (Kcal/mol) | DELTA G (Kcal/mol) | DELTA S (Cal/°K*mol) |
| --- | --- | --- | --- | --- | --- | --- | --- |
| Wild Type | 820804.2 | 51 | 85.7 | 85.7 | 22921.4 | 4302.8 | 60014.5 |
| DCIS | 815304.6 | 51 | 100 | 86 | 22765.4 | 4273.8 | 59604.8 |
|  | 817770.2 |  |  |  | 22832.1 | 4286.9 | 59777.5 |
| LCIS | 8177392 | 51 | 100 | 86 | 22838.8 | 4287 | 59799 |
|  | 8177739 | 52 |  |  | 22843.5 | 4290 | 59807.6 |
| DLCIS | 815304.6 | 51 | 100 | 86 | 22765.4 | 4273.8 | 59604.8 |
| LC | 810913.8 | 51 | 100 | 86 | 22633.7 | 4247.7 | 59264.3 |
|  | 828548.2 | 52 |  |  | 23137.9 | 4342.8 | 60583.7 |
| DC | 817466 | 51 | 99.5 | 85.7 | 22827.6 | 4286 | 59866 |
|  | 818960 |  | 99.9 | 85.9 | 22875 | 4294.6 | 59891.2 |

As an example biophysical properties are equally changed in the samples of DLCIS whereas, in LC, sample ID 992832 has got remarkable changes due to substitution type of mutation. The molecular weight has been increased from 820804.2 gm/mol to 828548.2 gm/mol. The enthalpy (ΔH), free energy (ΔG) and entropy (ΔS) are also increased from 22921.4(Kcal/mol), 4302.8 (Kcal/mol) and 60014.5 Cal/○K*mol to 23137.9 (Kcal/mol), 4342.8 (Kcal/mol) and 60583.7 Cal/○K*mol respectively. In LCIS sampleID 997326 the biophysical properties are decreased the most. The molecular weight has been decreased from 820804.2 gm/mol to 815000.4 gm/mol. The enthalpy (ΔH), free energy (ΔG) and entropy (ΔS) are also increased from 22921.4 (Kcal/mol), 4302.8 (Kcal/mol) and 60014.5 Cal/○K*mol to 22757.4 (Kcal/mol), 4272.6 (Kcal/mol) and 59582.9 Cal/○K*mol respectively.

### 3.2 Investigation of Protein at Sequential Level

As the function of a protein depends on its structure and structural information are embedded in the primary sequence, which is obtained from the translated nucleotide sequence; mutation of the DNA sequence will change the sequence of the expressed protein and may cause the sequences responsible for disease. The best example is the sickle cell anemia [15]. However, here in this present work, frame shifts deletion or insertion have made huge changes in the wild type DNA sequences resulting a large change in the primary sequence of the expressed proteins. Hence, we have extended our present work and tried to get an insight in sequential level of the expressed proteins.

#### 3.2.1 Understanding the trend of mutations in protein sequential level

In this subsection pair wise alignment of the expressed wild type and the mutated one has been done to identify the mutations in amino acid sequence. Table 6 shows some of such examples. The mutations change the codon sequences, as a result that lead to change in primary sequence of the proteins.

**Table 6:**
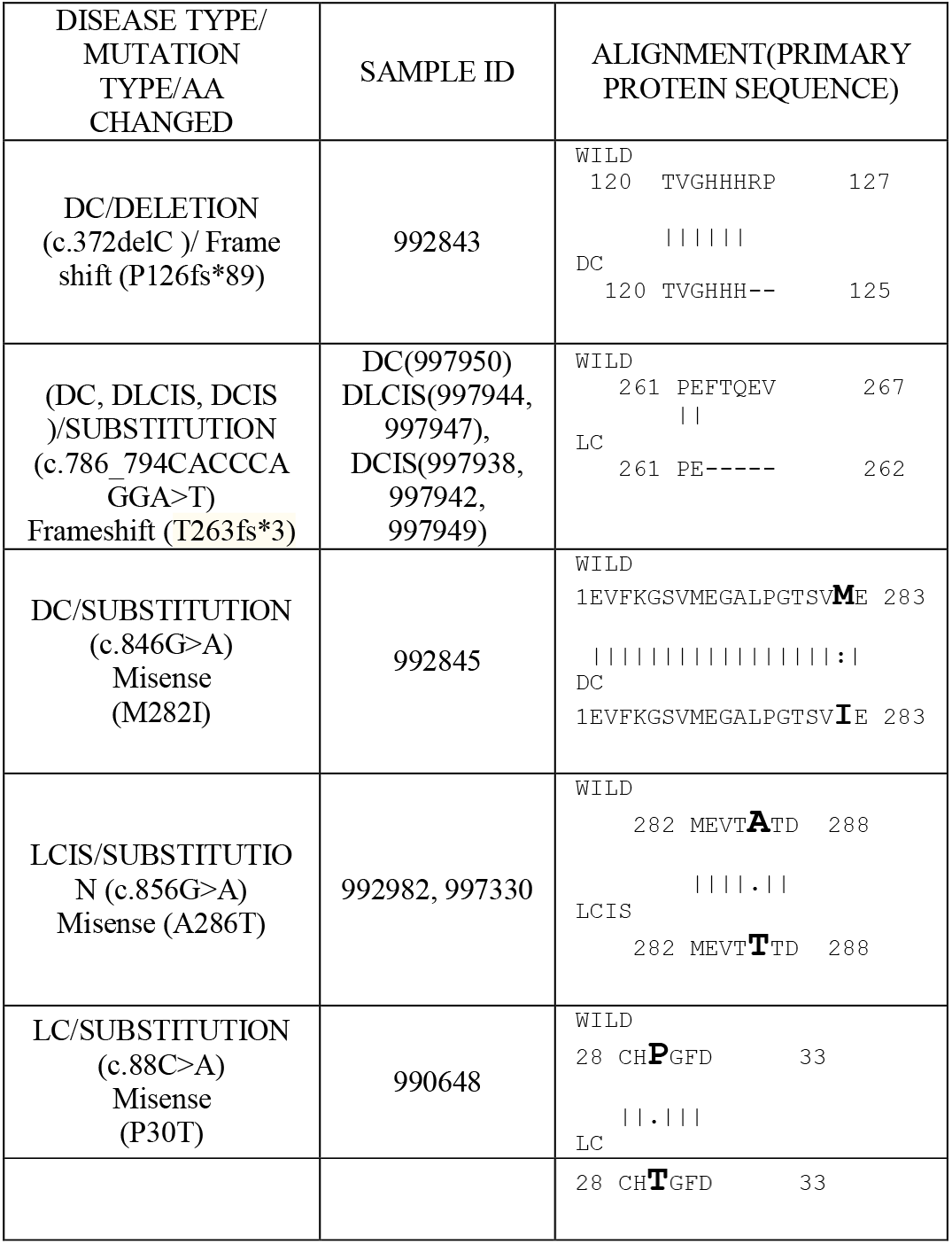
Example of pair wise alignments of normal and diseased proteins.

#### 3.2.2 Effect on Isoelectric Point of the mutated Protein Sequence

Isoelectric Point of a particular protein is a characteristic measure of its biophysical properties and helps to recognize where on a 2-D gel a particular protein can be found. In order to explore whether mutation in the wild type causes any change in their biophysical parameter(s), Isoelectric Point (pI) for all the mutated data have been derived along with wild type sequence using online software http://isoelectric.org/ (Supplementary table ST1). It can be observed from fig. 2 that the pI of the wild type CDH1 is 4.47 while due to mutations the isoelectric point of CDH1 protein has been changed in some cases. The pI of the mutated sequences responsible for LC (53 samples) show changes in the range of 4.47 to 11.02; whereas for those in DC (5 samples) it ranges between 4.47 to 10.83; for both DCIS (2 samples) and DLCIS (4 samples) suffer a huge change from 4.47 to 8.67.while sequence for LCIS it changes from 4.45 to 4.62. However, in DLC(1 sample) it is remained unchanged. This strongly recommend that due to change in pI the biophysical character of the mutated sequences get affected and resulting in disruption of the function, causing diseases.

**Fig 2:**
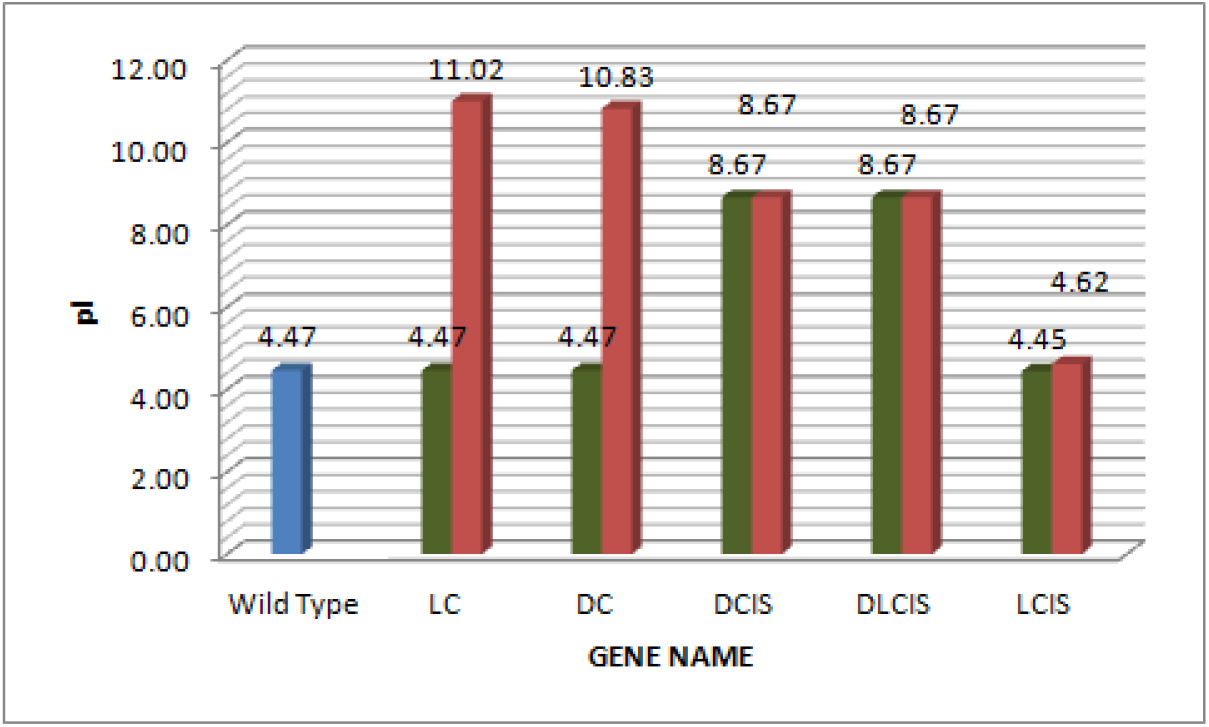
Determine Isoelectric Point(pI) of Protein Sequence. Here in this column chart the column of color blue represents the pI of wild type sequence. The green column represents the minimum range and the maroon columns represent maximum range of pI respectively for each type of breast cancer.

Towards better understanding of our proposition a case to case observation are cited. It has been observed that in the sample ID ‘992803’ (having LC type of breast cancer), the mutation is of deletion frame-shift type. 22 nucleotides are deleted between the CDS area 22 to 43. In sample ID ‘990647’ mutations take place due to insertion of 2 nucleotides A and G in the CDS area 49 to 50. In both the cases the isoelectric point (pI) is increased from 4.47 to 11.02. Sample ID 992843 having DC type of breast cancer affected by deletion type of mutation. At CDS area nucleotide C at 372 position is deleted and thus the isoelectronic point is changed to 10.83. DCIS type of mutations Sample ID(997938, 997942, 997949) witnessed substitution type of mutations (c.786_794CACCCAGGA>T) and is responsible for huge changes occurred in isoelectric point (8.67). Hence, the observation depicts that even a genotypic change in a single position in CDS area due to mutation can make huge changes in its isoelectric point and alter the normal function.

#### 3.2.3 Quantitative Analysis of severity and stability of mutation

Non-synonymous mutations lead to change in amino acids and cause structural changes of proteins. But all the mutations are neither harmful nor affects the biological functionalities of the protein equally. Online software PROVEAN (Protein Variation Effect Analyzer) v1.1 is used to find the virulence of a mutation, whether it is deleterious or neutral. If the PROVEAN score is ≤ a predefined threshold (e.g. - 2.5), the protein variant is predicted to have a “deleterious” effect else the variant is predicted to have a “neutral” effect [16–17]. Due to mutations Gibb’s free energy of the variants are changed, which affect in their stabilities. I-Mutant is used to predict protein stability changes due to single point mutations [18]. According to the methodology applied here in this online software, changes of protein stability (ΔΔ**G**) are measured in terms of Kcal/mol. ΔΔG (change of free energy of the mutant) <-0.5 indicates larger decrease in stability and ΔΔG > 0.5 indicates larger increase in stability. ΔΔG between −0.5 to 0.5 is considered as neutral stability.

Here in this subsection we have considered single point mutations and both the online software are used. It is observed that most of the mutations are deleterious (71%). On a keen look one can classify the effect of mutations into 4 states; Deleterious and stability decreased (54%), deleterious and stability increased (17%), neutral but stability decrease (25%) and neutral but stability increased(4%). From the results shown in Table 7 it can be concluded that due to mutation, stabilities are decreased (79 %) and hence biological functionalities of most of the variants are affected causing disease.

**Fig 3:**
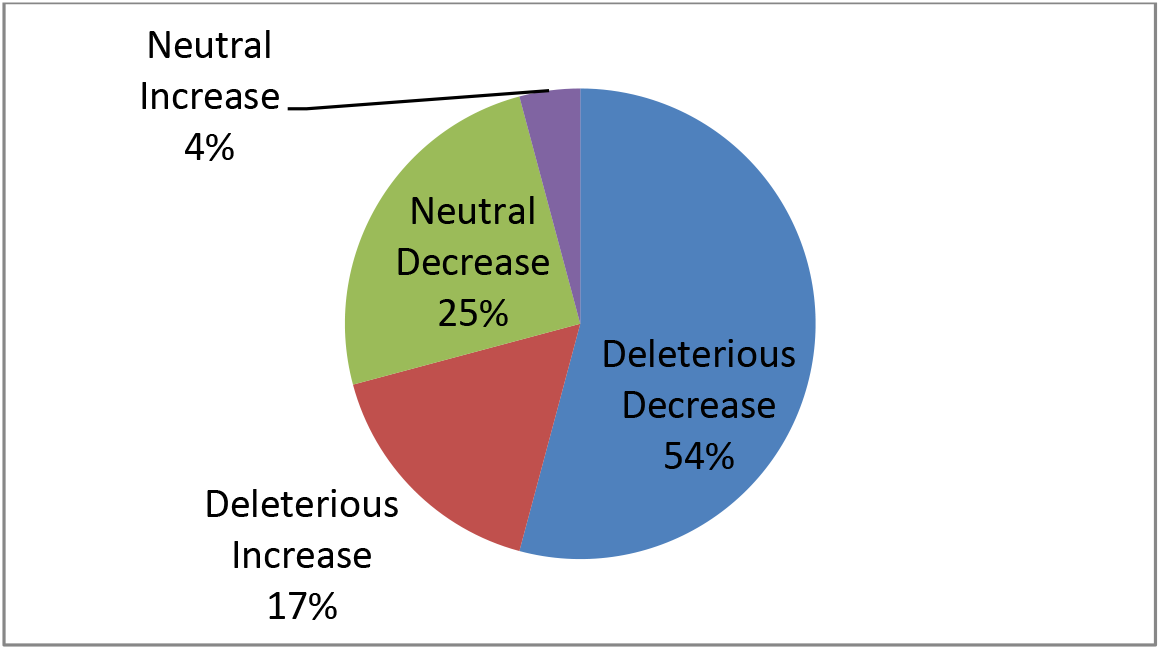
Prediction of biological functionalities changed due to mutations

**Table 7:**
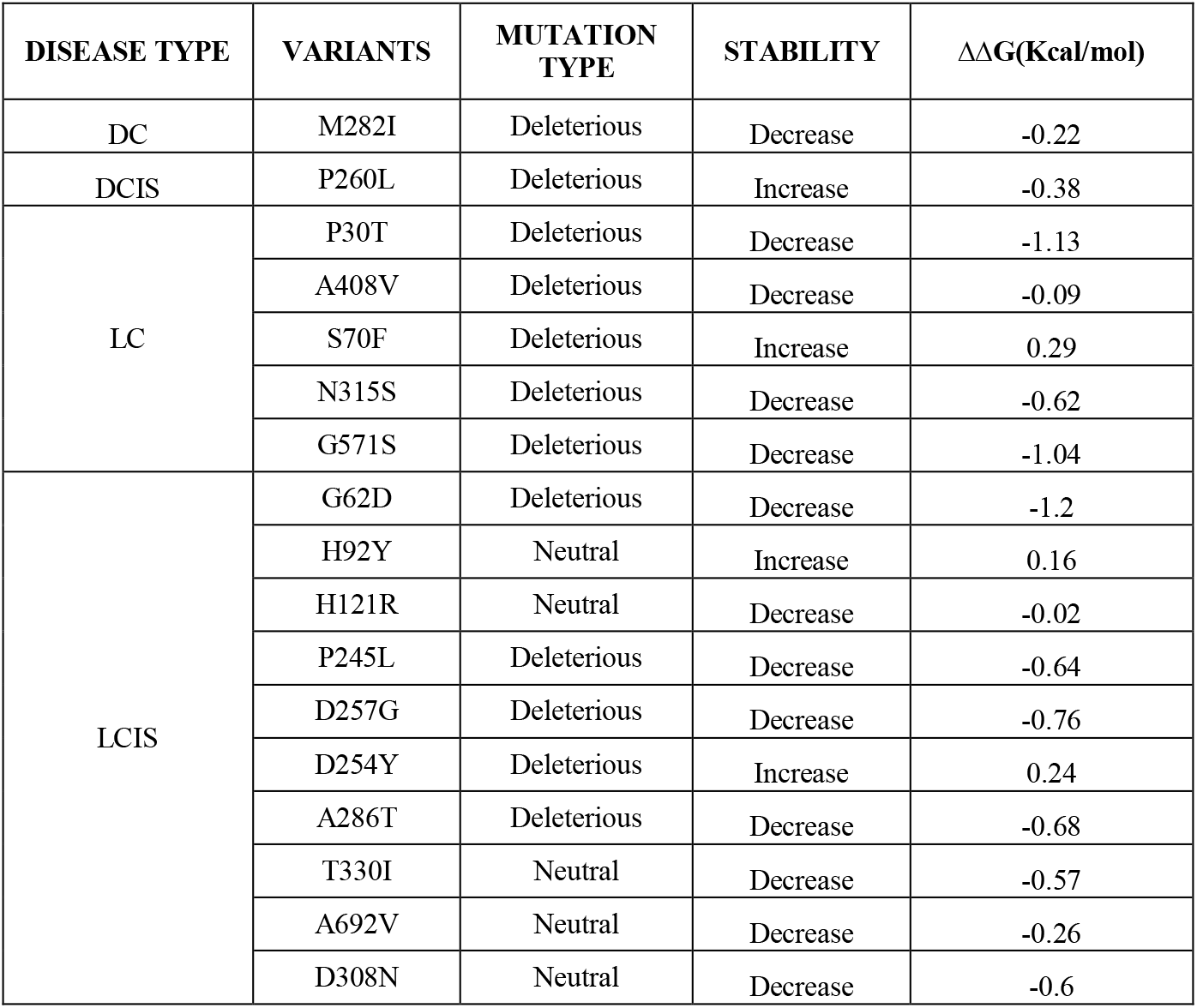
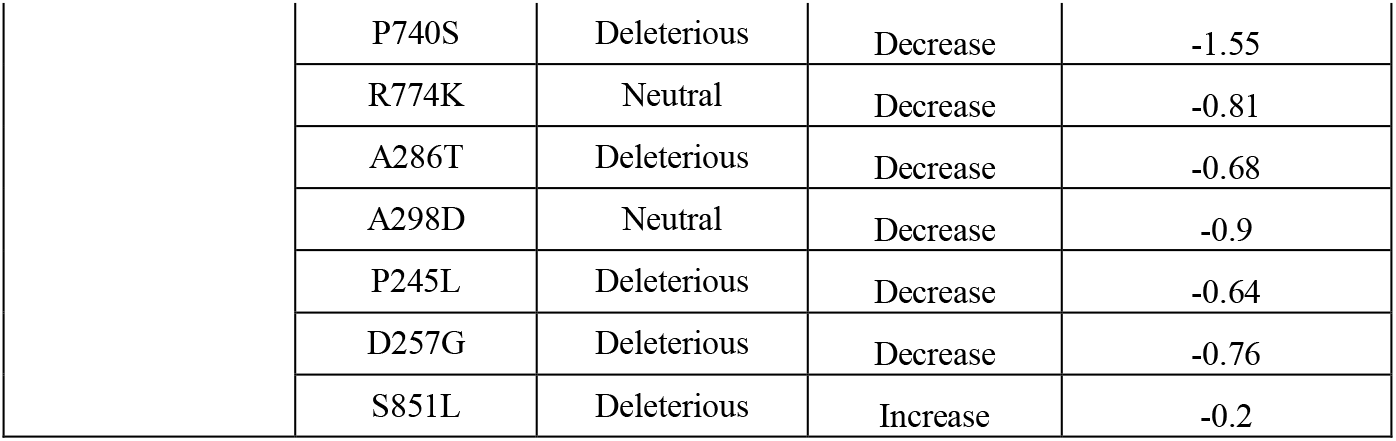
Quantitative Analysis of severity and stability of mutation.

## 4. Conclusion

Genetic information is stored in nucleotide sequences which through proper translation in expressed proteins maintain the proper biological functions. However, from the analysis of the patient database although not huge, one can easily hypothesize that mutations found in CDH1 gene, plays a significant role in breast carcinoma. As mature protein of CDH1 belongs to the family of cell–cell adhesion molecules, the mutations take places in different epithelial tissues of human breasts and thus participate in almost all types of breast malignancies. In this manuscript through identification of site of mutation in the CDH1 nucleotide sequence along with comparison and contrast between them with respect to the wild type one can correlate with the degree of pathology if possible. However, our study opens an avenue of the prediction of the protein secondary structures of the expressed mutated proteins which would give a better impetus towards understanding flaw in normal PPI and diagnose the rationality of breast cancer caused due to aberration of CDH1. In a nutshell, a consensus can be generated that woman with abnormal CDH1 gene (i.e. mutation) leads to have an increased risk of breast cancer.

